# Infection and host-feeding patterns of West Nile virus vectors varies by urban greenspace composition

**DOI:** 10.1101/2025.03.24.645034

**Authors:** Andrew J. Mackay, Jiayue Yan, Chang-Hyun Kim, Corrado Cara, Leta Chesser, Edna Alfaro, Michael Avara, Michael Ward, Seth Magle, Maureen Murray, Chris M. Stone

## Abstract

Greenspaces are integral to the urban environment and affect climate resilience, yet the extent to which they affect mosquito and West Nile virus (WNV) host populations and behavior is not clear. To address this question, we collected mosquitoes along a transect spanning a range of urban development. Mosquitoes were tested for West Nile virus, and the host species that were fed on determined for blood-fed specimens. Bird counts and camera traps were used to assess avian and mammal species availability. Different components that contribute to WNV transmission intensity responded to different landcover variables. Abundance of *Culex* mosquitoes was most strongly tied to impervious surface, while prevalence of infection was associated with increasing amounts of turf grass. The amount of turf was itself correlated with a measure of avian community reservoir competence. Blood meal analysis suggested the majority of blood meals in the ornithophagic species *Cx. pipiens* and *Cx. restuans* came from American robins and northern cardinals, with the latter in particular being overutilized relative to their abundance in sites with higher WNV prevalence. This work furthers our understanding of how the design of urban greenspaces could benefit from consideration of vector-host-virus interactions.

## Introduction

West Nile virus (WNV) is a zoonotic pathogen of considerable public health concern. It was initially isolated in 1937 in Uganda and originally occurred in Africa, the Mediterranean, East and Central Africa, with limited outbreaks in Europe, and generally thought to be of limited virulence [1]. Following the emergence of a virus strain of greater virulence, a number of larger outbreaks occurred, including in Romania, Russia, Tunisia, and Israel [1]. Following its introduction in 1999 in New York, it spread rapidly over the course of a 4-year-period across the continental United States [2]. West Nile virus is maintained in a cycle consisting of multiple avian hosts and mainly *Culex* spp. mosquitoes, and bird migration pathways are thought to have played a role in the rapid spread across northern America [3]. At the same time, it has affected bird populations and led to marked decreases in the abundance of several avian species [4]. While humans are considered dead-end hosts (i.e., incapable of sustaining a level of viremia that could lead to forward transmission), WNV can lead to considerable morbidity, with symptoms in neuroinvasive cases including meningitis and encephalitis, and these symptoms can sometimes lead to mortality [5]. Due to the animal reservoirs of this arbovirus, elimination is currently not feasible and public health responses are largely limited to preventive methods such as suppression of mosquito populations through vector control methods. How urban structure affects WNV transmission, and whether this could inform urban design in a way that mitigates the burden of WNV, have received relatively little attention.

West Nile virus incidence has frequently been tied to urban environments, in part due to the larger numbers of human cases that occur there as a result of high human population densities, and likely due to the ability of important vectors such as *Cx. pipiens* and *Cx. restuans* and key amplifying hosts (e.g., American robins, house sparrows) to thrive in urban environments [6,7]. These patterns however do vary regionally with the ecology of the primary vectors [6], and relative risk of exposure has been shown to vary from the absolute number of reported human cases as well [8]. The link to urban environments and vector-borne disease incidence is however concerning, as globally the proportion of humanity living in cities exceeds 50% and is expected to continue to increase [9].

At the same time, urban environments will need to adapt to climate change, with increasing global temperatures exacerbated by urban heat island effects [10]. One way in which urban planning could attempt to reduce the effects of urban heat islands is through nature-based solutions such as urban greening [11,12]. Urban green infrastructure and spaces of different types and compositions could affect mosquitoes or WNV transmission [13,14]. For instance, urban wetlands have been tied to lower WNV infection rates in mosquitoes [15,16], possibly due to effects on avian host communities or a negative association between *Cx. pipiens* and wetlands [17]. Vegetation density or composition in urban environments has also been linked to WNV incidence or *Culex* abundance [18,19]. Vegetation and green space could influence WNV transmission through a number of mechanisms. For instance, they could influence hydrology and the resulting availability and quality of larval developments sites for the primary WNV vectors. Other possible mechanism inlcude changes to habitat suitability for avian species with potential impacts on abundance and composition of avian communities; effects on local temperatures with ramifications for mosquito life histories and viral amplification rates; effects on predators [20] or mosquito microbiome composition [21], or by changing heterogeneity across the landscape in any of these factors.

Due to this multiplicity of potentially interacting factors, with effects of land cover, temperature, and host communities on mosquito production, feeding, and WNV prevalence, understanding how urban greenspace composition will affect WNV transmission remains challenging. The effects of temperature on the basic reproduction number of WNV illustrates this, as predictions are that transmission should be at its peak at a temperature of around 24-25 °C [22]. While the cooling effect of urban green spaces has potential to alter WNV transmission intensity, the magnitude and direction of these impacts might be expected to vary seasonally and in relation to latitude. Similarly, the effects of host community composition and diversity on WNV transmission can be scale- or context dependent, with certain studies highlighting effects of avian diversity or community competence on WNV prevalence [23,24], with others highlighting that foraging preferences for competent hosts suggest the importance of a few key reservoir hosts, such as American robins [25]. However, this can differ by region (e.g., [26]) and host selection patterns can vary throughout an urban core [27,28], although the reasons for this plasticity in host usage, relative to host abundance, remain poorly understood.

At the same time, *Culex* population dynamics may be influenced by environmental drivers in different ways [29]. For instance, the presence of water-holding containers, aging stormwater infrastructure, or other water bodies likely to hold nutrient-dense water for extended periods following precipitation events are thought to contribute to *Culex* abundance, yet it is not clear whether and how the environmental factors that optimize blood-feeding patterns differ from the way in which *Culex* abundance is maximized in terms of transmission.

An important question therefore is how mosquito and avian communities jointly are affected by urban environmental factors such as the composition and extent of urban greenspace and jointly shape important components or indicators of WNV transmission intensity. The main aims of this study were therefore to understand how factors related to transmission intensity, such as vector abundance, *Culex* spp. host feeding patterns, WNV infection rates as well as a composite measure of prevalence and vector abundance varied along a gradient of urban development intensity in the Greater Chicago area, and to assess these transmission-related parameters in relation to fine-scale urban land cover and host data.

We found distinct differences among sites in *Culex* species abundance and infection rates. Notably, abundance was correlated with environments high in impervious surface, while sites with higher WNV prevalence were positively correlated with the estimated extent of turf grass in the environment. The extent of local turf was correlated with avian community reservoir competence as a whole, and with a reservoir index for American robins and house sparrows. A comparison of the blood-feeding patterns and host usage in sites with varying infection levels suggested that northern cardinals were more strongly overutilized as hosts when they were rarer, and there was a weak relationship between the forage ratio on this species and WNV prevalence. Together, this suggests urban areas with high coverage of managed grass may favor avian communities and feeding patterns conducive to WNV transmission, which could inform urban design of natural spaces to promote health.

## Methods

### Study sites and mosquito sampling

Ten sampling sites were selected from locations previously utilized for a mammal diversity survey along a 50-km transect following the Chicago Sanitary and Ship Canal from the Chicago urban core to the southwestern suburbs [30]. Selected sites included urban parks, cemeteries, forest preserves and other natural areas, representing the diversity of housing/residential usage and natural/green areas within the Greater Chicago region (Fig 1).

**Fig. 1.**
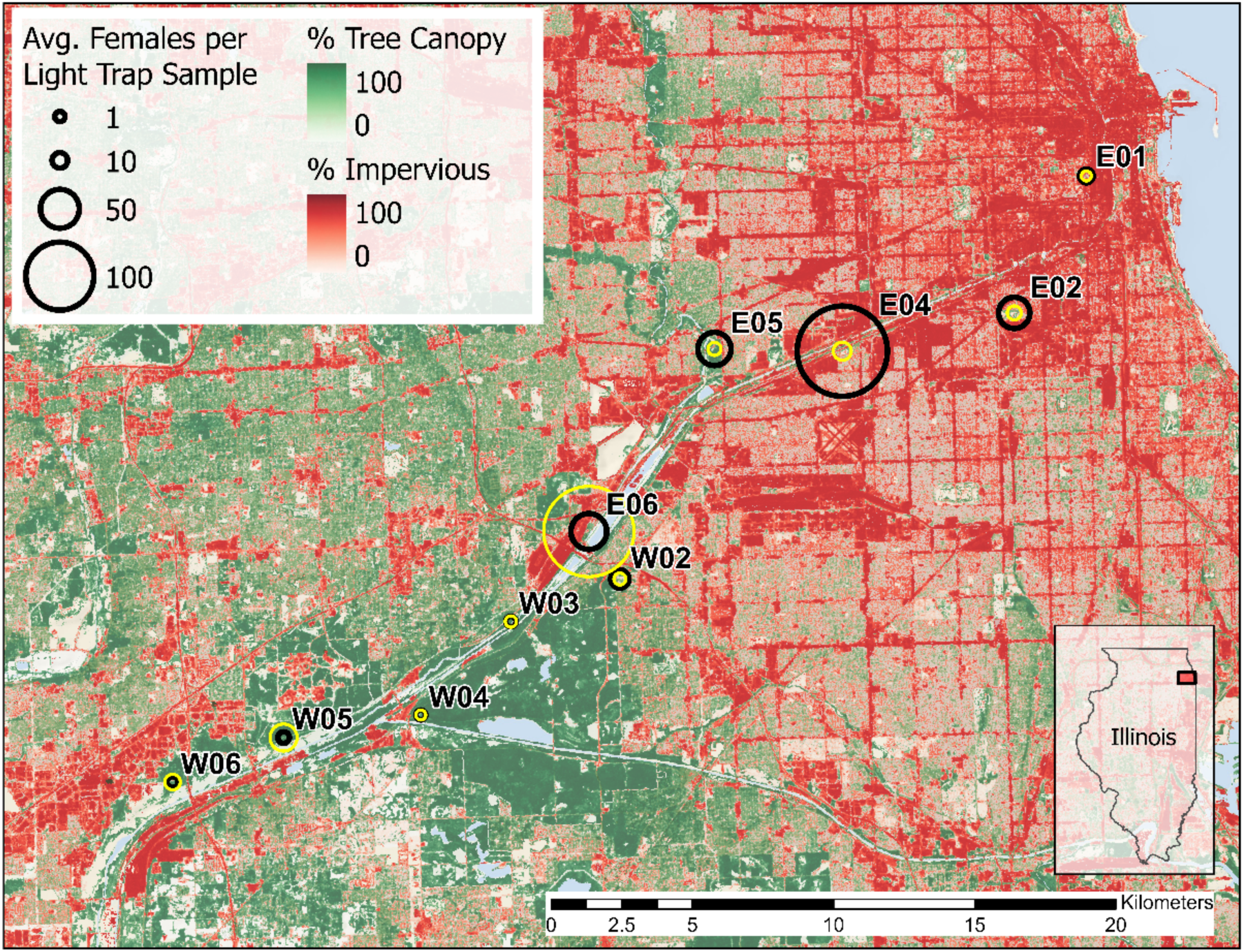
Land cover and the abundance of *Culex salinarius* (yellow) and all other *Culex* spp. female specimens (black; *Cx. pipiens*, *Cx. restuans* & unidentified *Cx.* spp.) in CDC light trap samples at ten sampling sites along a transect from urban (right) to suburban (left) areas in the Greater Chicago region

Mosquito sampling was conducted weekly in 2019 from June 25 to September 25 to coincide with the peak activity season of *Culex* mosquitoes and reported human WNV cases. Adult mosquito samples were collected weekly at each sampling site from four sampling locations per site spaced approximately 100 meters apart, except for the eastern-most site (i.e., the smallest natural area with the highest urban development intensity) where sampling was limited to three locations. Paired traps (min. 20 m apart) were placed at each sampling location within sites: a CDC gravid trap containing ∼6 liters of week-old grass infusion and a CDC light trap (light bulb removed) that was baited with ∼1.4 kg of dry ice and suspended at an approximate 1.5 m height. Additionally, a single BG Sentinel-2 trap (Biogents AG, Regensburg, German) baited with ∼1.4 kg of dry ice and a BG lure was used at each sampling site from 29 August 29 to September 25. For each weekly sampling event, traps were operated for an approximate 24-hour period, then samples were collected, transported to the lab on dry ice and stored at −80°C for subsequent identification and assays.

### Land Cover and Microclimate Measurements

Two primary data sources were used to characterize land cover within a 1000 m radius of our sampling sites: (i) continuous measures of canopy and impervious cover at a 30-meter resolution (NLCD 2019, [31]), and (ii) a 1-meter resolution dataset of classified urban land cover (MULC; [32]). Supplemental data were used to expand the original eight classes in the MULC dataset to ten land cover classes. First a building footprints layer [33] was used to separate impervious cover into “building impervious” and “non-building impervious” classes. To partition the grass / herbaceous cover class in the MULC dataset into “managed turf grass” and “non-turf” ground cover, the 2018 National Cropland Data Layer [34] was first used to identify cultivated agricultural crops possibly misclassified as grass / herbaceous cover. Potential misclassifications were subsequently compared with high resolution aerial imagery [35] for confirmation. A land use inventory [36] was then used to partition the proportion of remaining grass / herbaceous cover within land parcel boundaries into “managed turf grass” and “non-turf” grass / herbaceous cover, based on previously described turf grass relationships with parcel size, parcel land use and percent impervious cover ([37]; Table S1). For each site, average values of continuous NLCD impervious and canopy cover and the proportions of each MULC classified cover class were extracted within a 1000 m buffer radius from trap locations (ArcGIS Pro 2.7.1, ESRI, Redlands, CA) – a distance approximating the observed mean dispersal range of female *Culex* spp. mosquitoes in urban environments in the Chicago area [38].

To compare differences in microclimate across all sites, Thermochron iButton loggers (model DS1923-F5, Maxim Integrated, San Jose, CA, USA) were placed at a single, fixed CDC light trap location at each site to record air temperature and relative humidity at 10-minute intervals over a ∼24-hour period simultaneous with adult mosquito sampling, during a seven-week period from 23 July to 11 September. Loggers were suspended 30 cm from the light trap, at a height of ∼1.5 m, below an inverted 250 ml white paper cup to shield the logger from exposure to rain or direct sunlight. Each week, assignment of individual loggers to locations was randomized to minimize potential bias. Relative differences in air temperature and percent relative humidity for each 10-minute observation during the nocturnal period (sunset to sunrise) were calculated as the difference between the site value and global mean of values across all sites. Additionally, we calculated Vapor Pressure Deficit (VPD) for the four-hour period following sunset when adult *Culex* (*Cx*.) spp. are most active, and for the remainder of the nocturnal period when there is reduced flight activity [39–41].

### Species identification

The identification of female specimens to species was initially carried out on chill tables utilizing taxonomic keys [42–44]. For WNV screening, individuals lacking sufficient diagnostic characters to differentiate *Cx. pipiens*, *Cx. restuans*, and *Cx. salinarius* were pooled as *Culex (Cx.)* species. Of 198 blood-fed *Cx.* spp. specimens presenting morphological ambiguity, subsequent identification to species was achieved by multiplex PCR [45]. In summary, genomic DNA extracted for the blood meal analysis underwent multiplex PCR, targeting three forward primers specific to 28S ribosomal subunits: *Cx. pipiens*-specific PQ10 (5’-CCTATGTCCGCGTATACTA-3’), *Cx. restuans*-specific R6 (5’-CCAAACACCGGTACCCAA-3’), and *Cx. salinarius*-specific S20 (5’-TGAGAATACATACCACTGCT-3’), along with a common reverse primer CP16 (5’-GCGGGTACCATGCTTAAATTTAGGGGGTA-3’). Each 25-μl amplification reaction comprised 12.5-μl Q5^®^ Hot Start High-Fidelity 2X Master Mix (New England Biolabs, Ipswich, MA, USA), 3-μl forward primers (1-μl each), 1-μl reverse primer, 6.5-μl deionized water (dH_2_O), and 2-μl DNA sample. The PCR cycles were executed using Veriti^TM^ 96-Well Thermal Cyclers (Applied Biosystems, Waltham, MA, USA), involving one cycle at 96°C for 4 min, 40 cycles at 96°C for 30 sec, 51°C for 30 sec, 72°C for 90 sec, and one cycle at 72°C for 4 min. An equivalent volume (2 μl) of dH_2_O was included as a negative control for each PCR run. Verification of amplicons for *Cx. pipiens* (698 bp), *Cx. restuans* (506 bp), and *Cx. salinarius* (175 bp) was conducted through electrophoresis in 2% agarose gels at 180 V for 2 h 45 min in a large gel box. The identification of amplicon sizes relied on migration distance, with a 1-kb ladder (Invitrogen 1 kb plus; Thermo Fisher Scientific, Waltham, MA, USA) serving as a reference. Following molecular identification, 22 individuals that remained indistinguishable between *Cx. pipiens*, *Cx. restuans*, and *Cx. salinarius* were collectively categorized as *Cx.* spp.

### WNV infection

The WNV infection status of non-bloodfed *Cx.* spp. mosquitoes was determined by reverse transcription, real-time polymerase chain reaction (RT-qPCR) [46]. The QIAamp Virus BioRobot MDx kit (Qiagen, Germantown, MD) was used to isolate total RNA from pools containing a maximum of 50 specimens, aggregated by collection site, collection date and species. Five microliters of isolated RNA were utilized for a 20-microliter RT-qPCR reaction consisting of 1ξ TaqMan™ fast virus 1-step master mix (Thermo Fisher Scientific, Waltham, MA), 500 nM of each primer and 250 nM probe labeled with FAM fluorophore dye. Thermal cycling conditions included reverse transcription at 50 °C for 20 minutes, initial activation at 95 °C for 20 seconds followed by 40 cycles of PCR with a denaturation step at 95 °C for 30 seconds and an annealing/extending step at 60 °C for 60 seconds. The presence or absence of the WNV-specific gene in a given pool was determined from the amplification curve after setting a threshold line at 0.2. Mosquito pools with a cycle number at the threshold (Cq value) of 37 or less were considered positive for WNV RNA.

### Blood meal analysis

A total of 426 bloodfed *Culex* mosquitoes were processed individually for blood meal analysis. Genomic DNA of each individual specimen was isolated with NucleoSpin^®^ 96 DNA RapidLyse Kit (Macherey-Nagel, Allentown, PA, USA) following the manufacturer’s protocol. DNA concentration of each sample was measured using a dsDNA HS assay on a Qubit 3.0 fluorometer (Thermo Fisher Scientific, Waltham, MA, USA).

To mitigate the impact of PCR inhibitors present in the extracted DNA and account for the broad range of DNA concentrations (ranging from 0.1 to 120 ng/μl), a systematic dilution strategy was implemented. DNA samples exceeding 40 ng/μl underwent a 10-fold dilution, those within the range of 20-40 ng/μl underwent a 5-fold dilution, and samples below 20 ng/μl were used without dilution, with nuclease-free water as the diluent.

The diluted DNA samples were subjected to PCR using primers (forward: 5’-TNT TYT CMA CYA ACC ACA AAG A -3’; reverse: 5’-CAR AAG CTY ATG TTR TTY ATD CG -3’) targeting specific sequences of the vertebrate cytochrome c oxidase subunit I (COI) gene [47]. Each 25-μl amplification reaction consisted of 12.5-μl Q5^®^ Hot Start High-Fidelity 2X Master Mix (New England Biolabs, Ipswich, MA, USA), 1.25-μl of each forward and reverse primer, 7-μl dH_2_O, 1-μl dimethyl sulfoxide (DMSO), and 2-μl of the DNA sample. PCR cycles were conducted using Veriti^TM^ 96-Well Thermal Cyclers (Applied Biosystems, Waltham, MA, USA), involving denaturation at 94°C for 5 min, followed by 40 cycles of denaturation at 94°C for 40 sec, annealing at 55°C for 30 sec, extension at 72°C for 60 sec, and a final extension at 72°C for 7 min. Negative controls containing 2-μl dH_2_O were included in each PCR run.

Following amplification, PCR products (approximately 300kb) were visualized through electrophoresis in 2% agarose gels using the E-Gel^®^ Imager (Life Technologies, Carlsbad, CA, USA). Subsequently, all samples underwent size selection using PippinHT (Sage Science, Beverly, MA, USA) to isolate amplicons falling within the 280-320 kb range. The selected PCR products were purified using the Mag-Bind^®^ TotalPure NGS Kit (Omega Bio-Tek, Norcross, GA, USA), and their DNA concentrations were reassessed.

The purified samples then underwent an overhang PCR step to extend the primers with identification (ID) sequences. Following purification and concentration measurement, each sample was adjusted to 20 ng in a total volume of 50-μl for a third-round PCR. The third-round PCR utilized 25-μl Q5^®^ Hot Start High-Fidelity 2X Master Mix, 4-μl Nextera UD Indexes, 16-μl dH_2_O, and 5-μl of the sample. The indexed PCR was performed with an initial denaturation step at 95°C for 3 min, followed by 12 cycles of denaturation at 95°C for 30 sec, annealing at 55°C for 30 sec, extension at 72°C for 30 sec, and a final extension at 72°C for 5 min. After purification and concentration determination, 50 ng of each sample’s product was allocated to build two sequencing pools, which were then submitted for Next-Generation Sequencing (NGS) using the Illumina MiSeq Nano platform at the Edward R. Madigan Laboratory, University of Illinois at Urbana-Champaign.

The paired-end sequencing data underwent primer sequence removal utilizing cutadapt version 4.1 [48], employing default parameters, which encompassed a maximum error rate of 10% and utilization of the -g flag to eliminate bases situated upstream of the primers.

Subsequent amplicon processing was carried out in R software version 4.3.1 with the DADA2 package version 1.28.0 [49]. Read quality assessment was performed using the plotQualityProfile function. Filtering and trimming of low-quality reads were executed by adjusting parameters such as truncLen (225, 220), minLen (20), truncQ (2), and maxEE (2, 2) within the filterAndTrim function, while considering the overall sequence quality. Forward and reverse reads were merged using the mergePairs function, adhering to default parameters with a minimum overlap of 12 and no allowed mismatches. Chimeric sequences were subsequently eliminated using removeBimeraDenovo with default settings. The origin of each edited sequence was pinpointed through a BLAST search in the GenBank database. The top 1 hit, boasting a 98% identity and typically covering the entirety of the sequence (100%), was designated as the host species for each sample sequence. The species with the greatest number of reads associated with it for a given blood-fed specimen was designated as the host that mosquito fed on.

### Bird surveys and camera traps

Point counts were performed at each location at three dates between mid-July to mid-September for a 15-min period at each location. Counts were performed during the morning (on days without rain) between sunrise-10:30 am and the order in which sites were visited was randomized among the repeat visits. Counts were based on both visual (using binoculars) and auditory cues. Mean numbers of observations over the successive visits were calculated for each site.

Camera traps were set in the same urban greenspaces as mosquito collections occurred and were deployed for 30 successive days in July and October of 2019. Camera traps were operated and data collected as previously described [30].

### Statistical analysis

Analyses were performed using R version 4.4.1. We derived maximum likelihood estimates of the prevalence of infection for West Nile virus in mosquito pools of varying sizes using the PoolTestR package [50]. Correlation matrices were created using the PerformanceAnalytics package [51]. Monte Carlo multinomial tests were performed using the EMT package [52]. Linear regression analyses were used to assess land cover relationships with the abundance of host seeking females (average *Culex* spp, females collected per light trap sample) and maximum likelihood estimates of WNV infection rate (*Culex* spp, females pooled among all three collection methods), restricted to sampling dates when collections were made at all ten study sites. Model selection was performed using the stepAIC function of the MASS package [53].

## Results

### Abundance of Culex species mosquitoes

The overall numbers of collected female West Nile virus vector *Culex* (*Cx*.) spp. are given in Table 1. This includes the primary vectors *Cx. pipiens* and *Cx. restuans* as well as the possible bridge vector *Cx. salinarius*. There was considerable variation in the average numbers of female *Culex* spp. collected among the different sites, with generally a lower abundance in the sites along the western arm of the transect, compared to the sites closer to the urban core (Fig. 1). Exploring the relationship between female (log-transformed) abundance and urban land cover with linear regression analysis shows a significant positive correlation between the proportion of impervious surface within a 1000m radius surrounding the trap locations and *Cx. pipiens* abundance (*R* = 0.72. p = 0.018) as well as with *Cx. restuans* abundance (*R* = 0.71, p = 0.021). Conversely, the abundance declines with increasing proportions of land cover consisting of tree canopy, for both *Cx. pipiens* (*R* = −0.76, p = 0.010) and *Cx. restuans* (*R* = −0.67, p = 0.033) (Fig. 2).

**Fig. 2.**
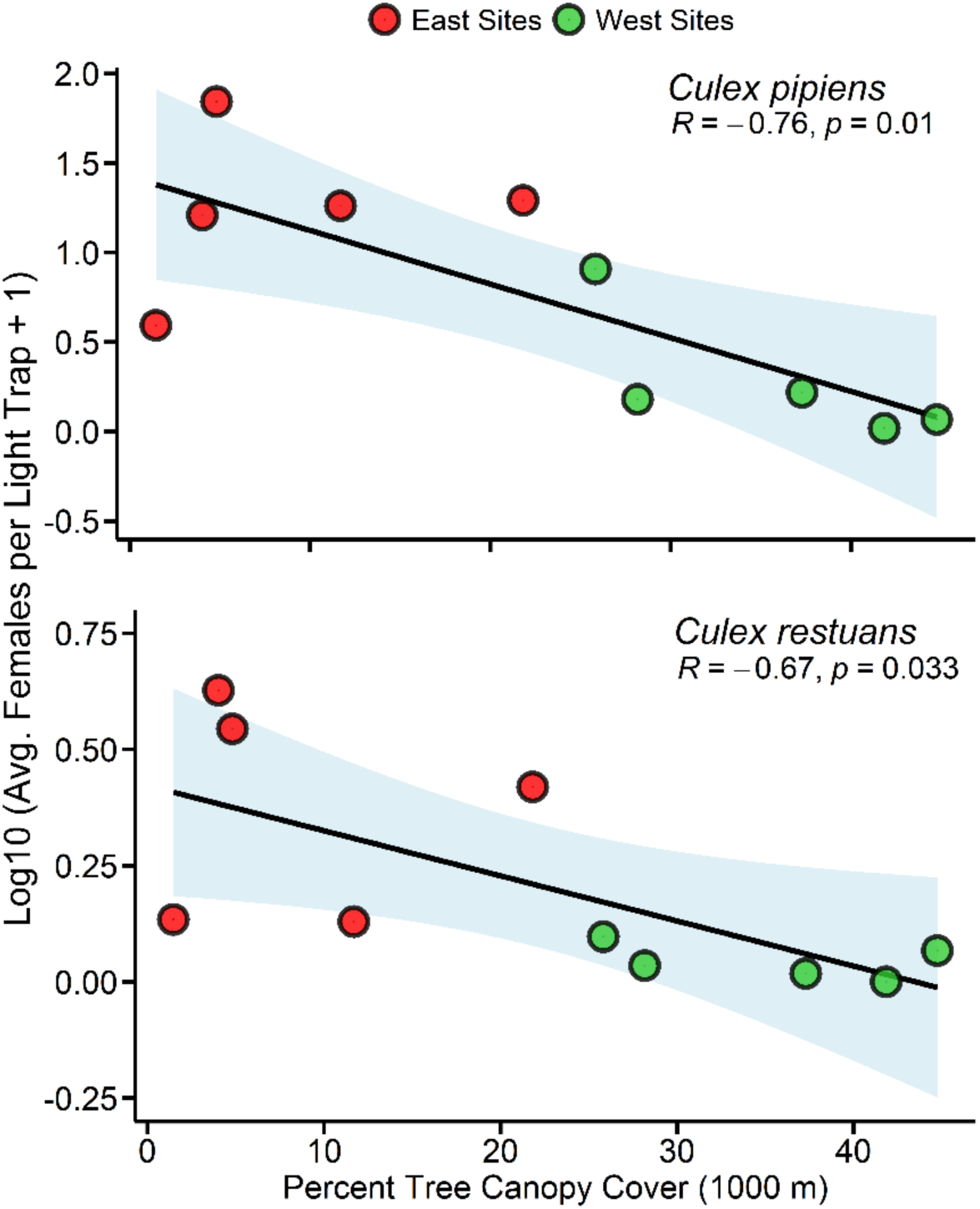
Average log-transformed abundance per site of *Cx. pipiens* and *Cx. restuans* females in CDC light trap samples by the percentage of tree canopy present in a 1000 m buffer around each site.

**Table 1:** Maximum likelihood estimates of West Nile virus infection rates by species.

| Species | WNV infection rate |  | Females | Pools |  |
| --- | --- | --- | --- | --- | --- |
|  | MLE | 95 % CI |  | Tested | WNV+ |
| *Culex (Cx.)* spp. | 5.5 | 4.0, 7.3 | 8,944 | 324 | 44 |
| *Culex pipiens* | 2.3 | 1.3, 3.7 | 6,377 | 255 | 14 |
| *Culex restuans* | 1.6 | 0.4, 4.2 | 1,930 | 184 | 3 |
| *Culex salinarius* | 0.2 | 0, 0.9 | 5,502 | 185 | 1 |
| *Culex tarsalis* | 0 | 0, 62 | 45 | 15 | 0 |
| **Total** |  |  | **22,798** | **963** | **62** |

For *Cx. salinarius*, which was only collected from light traps, there were no significant correlations with impervious surface. Both these points – the lack of attracting oviposition-site seeking females of this species in gravid traps and the lack of association with impervious surface likely result from the use of different types of larval development sites than what is typically associated with more urban environments. Alternatively, it could be that there are differences in the responses of these species to environmental factors like temperature and relative humidity, which we measured at each site and found to be correlated with canopy cover and impervious surface as well (Fig. S1).

### West Nile virus infection rates

An overview of the number of mosquito pools tested and the WNV infection rates per species is provided in Table 1. The maximum likelihood estimates of prevalence varied among the species. We observed a higher infection prevalence in *Cx*. spp., i.e., specimens that could not confidently be identified to species based on morphology. This most likely is because females in poor condition that have lost scales will tend to be older and therefore will have had more opportunity to bite and become infected. The rate of infection in *Cx. pipiens* was slightly higher than that of *Cx. restuans*, but with overlapping confidence intervals. The rate of infection in *Cx. salinarius* was considerably lower.

The prevalence of infection for all species varied among collection sites (Fig. 3a), in a manner that is distinct from the pattern observed for *Culex* abundance. Notably, a number of sites with less intensive urbanization saw relatively high levels of infection. When examining correlations with land cover, the strongest relationship was found with the proportion of land classified as turf grass (*R* = 0.78, p < 0.001). This suggests that the prevalence of infection is not as strongly coupled to mosquito abundance as to other factors, possibly to do with the composition of the avian host community.

**Fig. 3.**
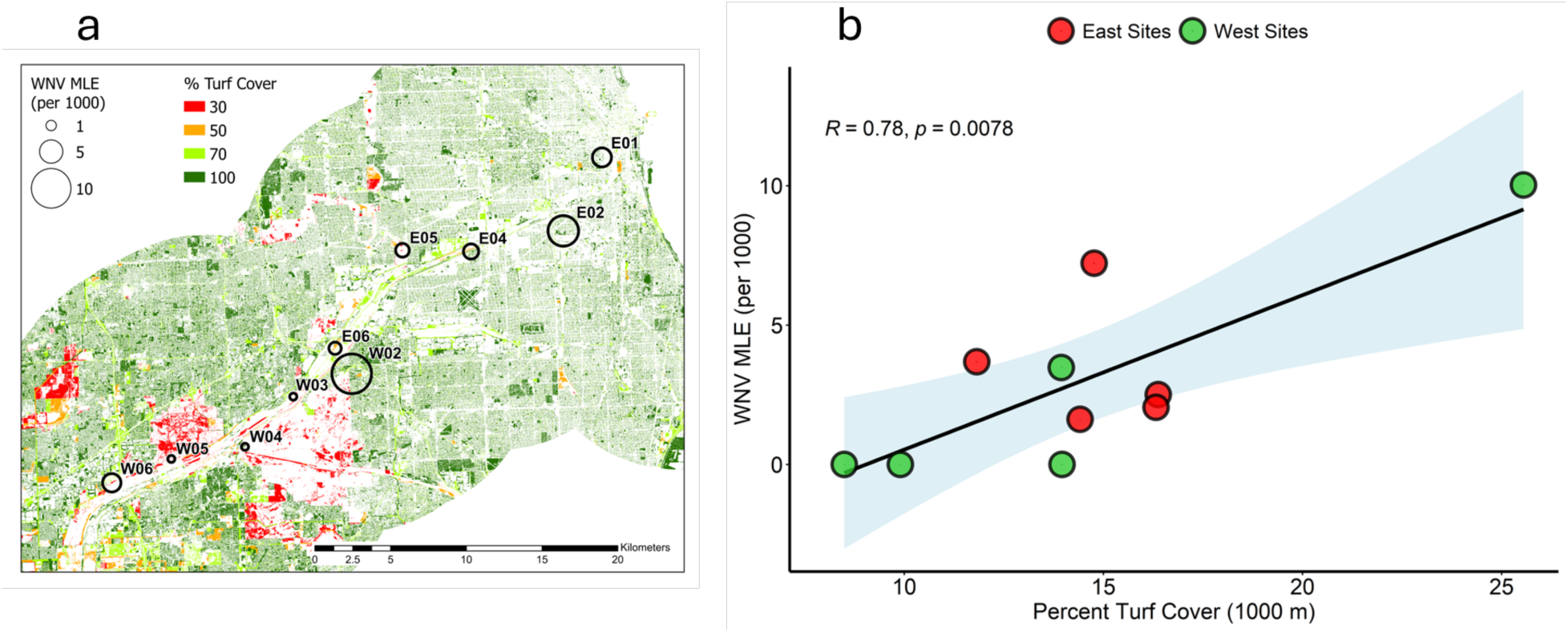
A) Maximum likelihood estimate of WNV prevalence in *Culex* spp. per site and the distribution of grass / herbaceous land cover, classified according to the percentage represented by turf grass. B) WNV prevalence as a function of the percentage of land area classified as turf grass cover in a 1000 m radius around collection sites.

Our avian point counts documented a total of 59 bird species. Overall, American robins (*Turdus migratorius*) were the most common species and their relative abundance appeared similar along the eastern and the western side of the transect. A number of other species appeared to differ in this regard, with European starlings (*Sturnus vulgaris*) and house sparrows (*Passer domesticus*) being more abundant in the more eastern sites, while blue jays and American goldfinches (*Spinus tristis*) were more common in the western sites (Fig. S2).

To explore whether and how WNV prevalence is influenced by avian species assemblages, we calculated a community reservoir competence. To do so we calculated for each avian species a competence index, *C_i_*, as the product of their reservoir competence, *c_i_*, based on values from the literature [54–56] and *a_i_*, the abundance of that host species at a given site. The sum of the *C_i_* values for all species observed at each site provided the community competence index (*CCi*) [23]. An overview of the variation among sites in this metric and the contributions of individual species to it is presented (Fig. 4). To explore the effect of the species’ *C_i_* values and environmental parameters on WNV prevalence, following univariate analyses and examining Pearson correlation plots, we performed a linear regression with WNV prevalence as the dependent variable and the *C_i_* values for American Goldfinch (*C_AMGO_*), the *CCi*, and the proportion of turf around the collection sites as predictor variables. The best model following stepwise selection based on BIC values included *C_AMGO_* (t=3.78. p = 0.007), and the proportion turf (t=2.87, p=0.026). It is possible that the proportion of land cover designated as turf serves as a proxy for other parameters more directly affecting WNV transmission. For instance, proportion turf cover was significantly positively correlated with *CCi*, and the *Ci* values for house sparrows and American robins, and in a univariate screen, *CCi* also was significantly correlated with WNV prevalence in mosquitoes (Fig. 4).

**Fig. 4.**
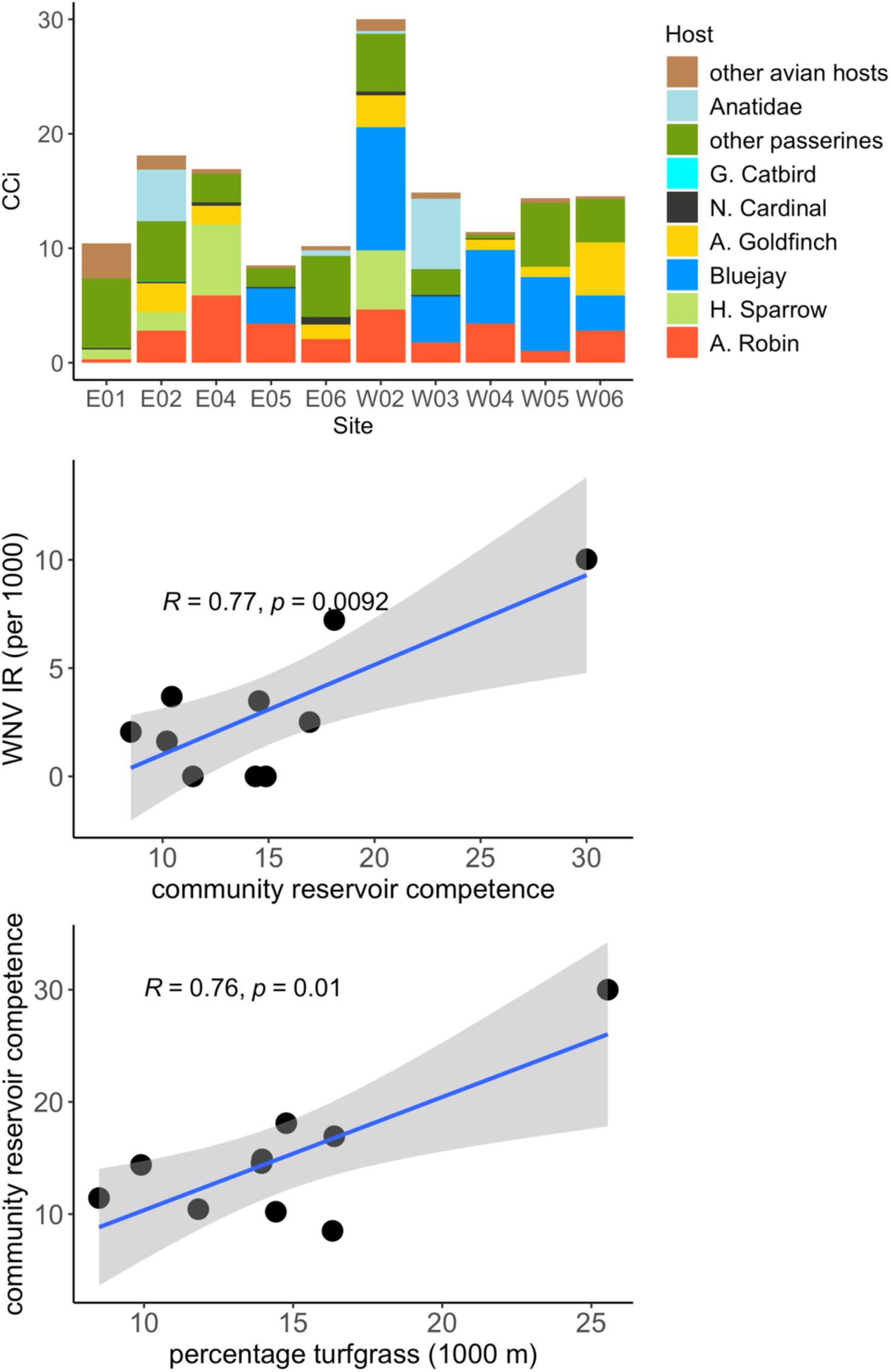
A) The contribution of avian species to the community competence index (CCI) for West Nile virus at different sites; B) The relationship between CCI and WNV infection rate in *Culex* spp.; C) The relationship between CCI and the proportion of turfgrass in a 1000m radius around each site.

### Blood feeding patterns and preferences

When looking at the relative frequency at which different vertebrate species were used as blood-hosts by the *Culex* species, it is evident that there are a relatively small number of species that are fed on at high rates, with a long tail of species that are fed on more infrequently (Fig. 5). The hosts used by *Cx. pipiens* and *Cx. restuans* appear to largely overlap. For *Cx*. *pipiens*, the most common hosts were American robins (24.2%), northern cardinals (*Cardinalis cardinalis*, 18.9%), gray catbirds (*Dumetella carolinensis*, 16.7%), and house sparrows (5.3%). The proportion of blood meals that came from humans was 3.8%.

**Fig. 5.**
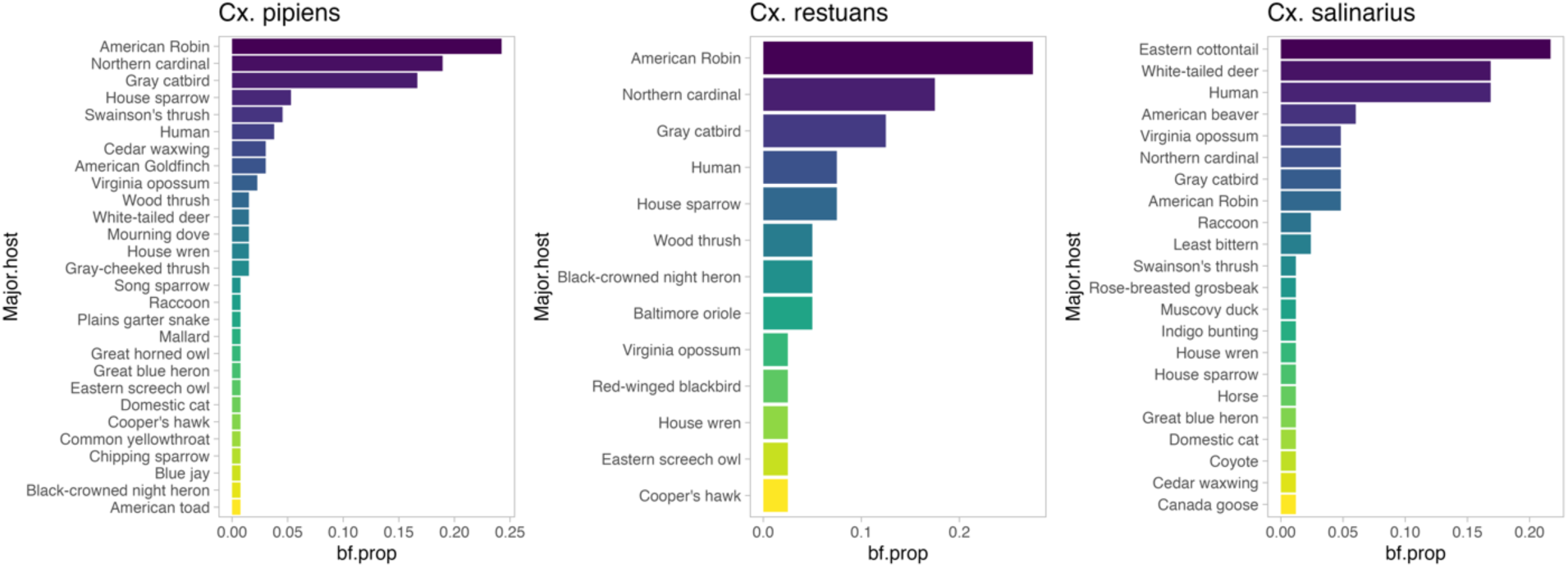
The proportion of blood meals taken on different vertebrate host species for *Cx. pipiens*, *Cx. restuans*, and *Cx. salinarius* across the sampling transect.

**Fig. 6.**
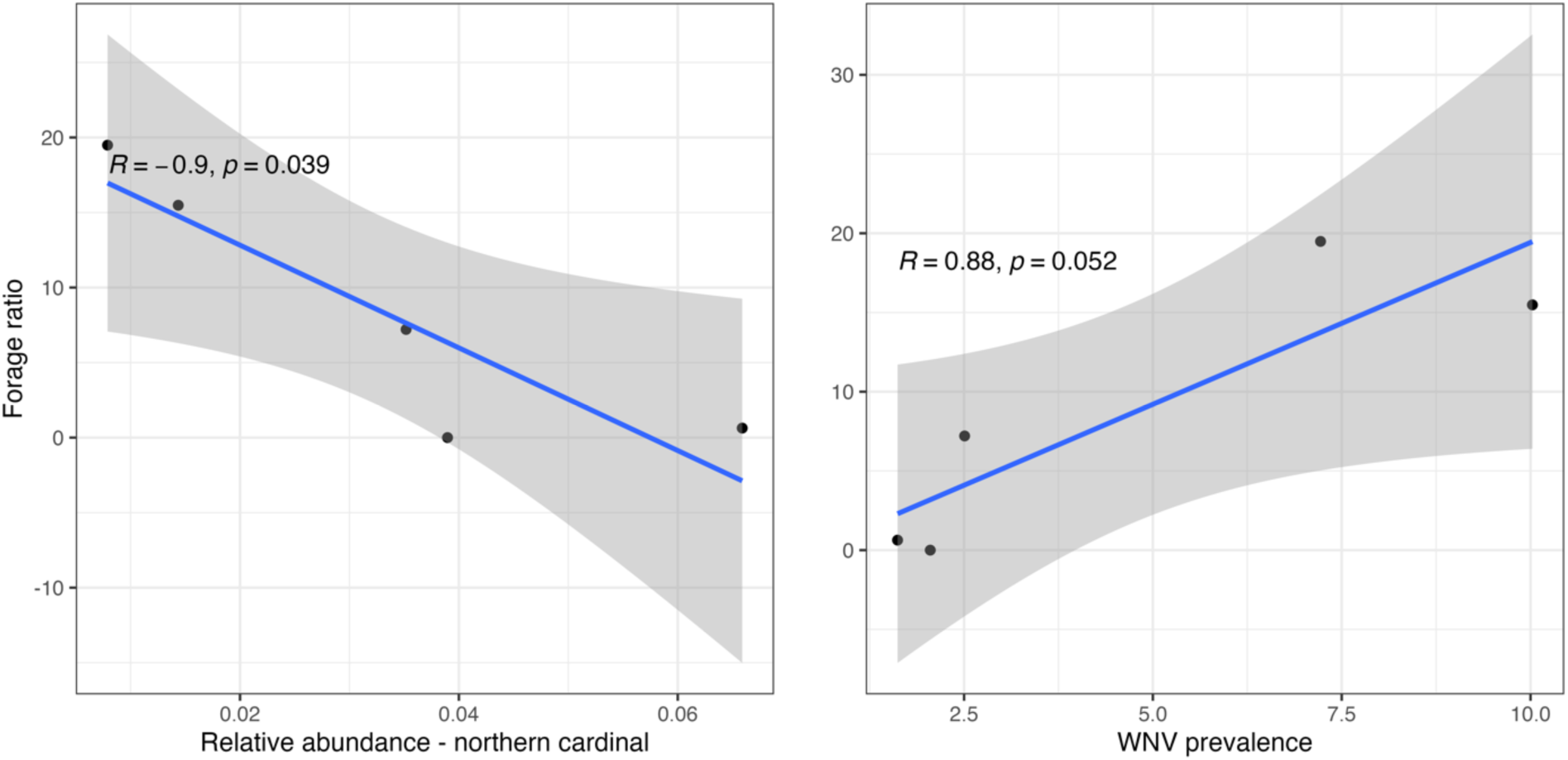
Left: the forage ratio on northern cardinals by *Cx. pipiens* as a function of the relative abundance of northern cardinals at different sites; Right: the forage ratio in relation to the WNV prevalence in *Culex pipiens* group mosquitoes at those sites.

We then compared the proportions of blood meals per host type among areas with the highest WNV infection prevalences and those with the lowest prevalences (Fig. S3). There was a significant difference in the distribution of blood meals among these environments (X=17.51, p=0.022). To contrast whether there were differences for individual species among the habitat types, we assessed whether the standardized residuals from a chi-square analysis had an absolute value greater than 1.96. Notably, this suggested that gray catbirds and American robins made up a larger proportion of blood meals in the lower prevalence sites. Other avian species (a number of species with only 1 or 2 blood meals each grouped together) made up a larger proportion of blood meals among *Cx. pipiens* in sites with higher infection rates. No significant differences in the proportion of blood meals between low and high infection sites were observed for any other host species or host group.

In addition to the proportion of blood meals taken on different hosts, host preference, or the rate at which different species are fed on relative to their availability, can also provide meaningful insights into infection dynamics. To explore this, we calculated forage ratios for some of the more commonly fed-on species as the proportion of blood meals on a species divided by its relative abundance. For American robins, the forage ratio was comparable among sites with high or low WNV infection prevalence (high: 1.21; low: 1.15). However, the forage ratios for house sparrows (high: 3.69; low: 0.98) and northern cardinals (high: 22.47; low: 10.13) strongly favored the high prevalence sites. For northern cardinals we saw a significant correlation (R=-0.9, p = 0.039) between the forage ratio and relative abundance of cardinals, so that as cardinals were rarer at a given site, the forage ratio on them increased, suggesting a relatively inelastic feeding rate on this species. There was also a trend for a relationship between WNV prevalence in primarily ornithophagic vectors (i.e., all *Culex* (*Cx.*) spp. excluding *Cx. salinarius*) and the forage ratio on cardinals, however, this was not significant (R = 0.88, p = 0.052).

The blood-feeding pattern we observed for *Cx. salinarius* differed considerably from the other two *Culex* species, with a majority of blood meals (72%) coming from mammals, primarily white-tailed deer (*Odocoileus virginianus*), eastern cottontails (*Sylvilagus floridanus*), and humans. Among birds, American robins, gray catbirds, and northern cardinals were most commonly fed on. When grouping blood-fed females by region (sites located in the eastern or western half of the transect), the most noticeable shift in blood feeding patterns related to a higher proportion of blood meals taken from white-tailed deer in the western region, with a greater proportion of females feeding on eastern cottontails and humans in the eastern region. These differences, however, were not statistically significant (Monte Carlo multinomial, p=0.09). The rate of detection (the number of days where this species was observed with a camera trap out of all the days the camera trap at a given location was active, averaged over July and September recordings) for white-tailed deer varied per site, with sites closer to the urban core (E01, E04), as well as W02, not recording any deer activity, and moderate to high rates at the other locations. Eastern cottontails on the other hand were only recorded on camera traps in the eastern sites, which could either reflect a difference in abundance, observability, or a combination of both. A large proportion of blood-fed *Cx. salinarius* came from one location (E06). Here, white-tailed deer had a detection rate of 71.5% and eastern cottontails one of 18.5%, while 5 and 15 females bloodfed on these species, respectively. A feeding index [57], calculated as 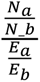, with *N_i_* representing the number of blood meals on species *i*, and *E_i_* the expectation of the numbers of blood meals based on their relative abundance, based on these two species at this site is then 0.09, which suggests a relative underutilization of white-tailed deer. The number of observed bites on white-tailed deer at this site over eastern cottontails differs significantly from the expected numbers based on their average rates of detection by the camera trap (χ^2^=35.2, p < 0.05).

## Discussion

Environmental factors influence vector-borne disease transmission through complex interactions between vectors, host communities, and pathogens, each of which may be affected differently by the same environmental condition. Understanding these ecological interactions allows for a better insight into areas where people may be most at risk, and potentially could lead to insights into how the environment could be managed in order to decrease transmission of pathogens such as WNV. Here, we explored how variation in urban land cover affects host and vector communities and how these jointly affect WNV vector infection rates. We found that there was a discrepancy between the factors influencing vector abundance and infection rates, with *Culex* species abundance being positively correlated with increasing urban development and impervious surfaces, and WNV prevalence being positively associated with greater turf grass cover. The amount of estimated turf grass was also strongly correlated with the community reservoir competence of resident avian host communities as well as the availability of certain important reservoir hosts, such as American robins.

The most common avian hosts that were fed on were American robins, northern cardinals, and gray catbirds. American robins being the most commonly fed-on host is consistent with other studies on *Culex* blood-feeding patterns in the northeastern parts of the U.S [25,27,58]. In our field sites, American robins were also the most commonly-observed species overall, suggesting *Culex* spp. in this region are not overusing this species in the same way as has been reported in other locations [25]. Given their abundance and the high proportion of blood meals being taken on this species, American robins clearly contribute strongly to the force of infection on mosquitoes in this region. The competence index, *C_i_*, for robins was also correlated with the overall community reservoir competence, and with the amount of estimated turf grass around collection sites. However, neither estimates of the competence index of robins alone nor the estimated forage ratio on robins were significantly associated with WNV prevalence. This suggests that American robins are an important contributor to community reservoir competence, but not on their own sufficient to explain patterns in WNV prevalence in this study.

We also found that both *Cx. pipiens* and *Cx. restuans* fed on humans a relatively small proportion of the time (3.8% and 7.5%, respectively), which supports the notion of these species serving both as primary amplification vectors and as bridge vectors leading to spillover to humans [59]. Observed blood-feeding patterns of *Cx. salinarius* and its’ relatively high competence for transmitting WNV have led to speculation to its putative importance as a bridge vector in certain areas [58,60,61]. In our sites, this species fed on humans to a greater degree than *Cx. pipiens* and *Cx. restuans*, while also feeding on avian species such as American robins and northern cardinals, which supports the possibility of this species serving as a bridge vector. This will however be tempered by their lower abundance, heterogeneous distribution, and the much lower level of WNV prevalence in this species. A notable finding was that the most common host for this species was the eastern cottontail, with white-tailed deer being second, and a feeding preference for cottontails observed at a site where both species occurred. In the more wooded western areas, white-tailed deer were the most commonly fed-upon species. Whether either of these two blood-hosts could effectively divert this species from feeding on humans (i.e., zooprophylaxis) is not something we were able to ascertain here, due to the lack of information on human occurrences at these locations, but it warrants further research.

Mosquitoes in the high infection rate sites were more likely to have fed on avian species represented by only 1 or 2 blood meals (i.e., “other Avian” group), which included such species as the American goldfinch, black-crowned night heron (*Nycticorax nycticorax*), and the common yellowthroat (*Geothlypis trichas*). While a significant proportion of blood meals came from American robins, surprisingly the proportion of robin was higher in the sites with low infection prevalence (Fig. S3). Notably, we observed a greater degree of feeding on gray catbirds in the low infection prevalence sites. In Georgia, a high rate of feeding on mimid species was found and was suggested to lead to a dampening of transmission due to their low competence index for WNV [26], and it is possible such a dampening effect occurs in certain sites in the Greater Chicago area as well.

A further difference in feeding patterns that could contribute to the observed higher prevalence levels at certain sites is the greater forage ratio on certain WNV hosts, namely house sparrows and, in particular, northern cardinals. The forage ratio represents a measure of the extent to which a particular species is under- or overutilized relative to its abundance. The role of cardinals in WNV transmission remains somewhat unclear due to their moderate competence [26]. It may be that the strong preference for cardinals in certain locations, along with the resulting heterogeneity in feeding, makes up for this and leads to an overall increase in transmission intensity [62]. In Louisiana, cardinals appeared to be among the primary amplification reservoir species for WNV [63,64] and experimental infection studies show that they sustain high levels of viremia capable of infecting mosquitoes [64]. Additionally, northern cardinals have in a number of studies been reported as having some of the highest seroprevalence rates, including in Illinois [65,66], likely resulting from the biting preferences of *Culex* species. A significant concentration of biting on this species could thus plausibly be involved in intensifying transmission in certain environments.

An area in need of further research is understanding what leads to changes in feeding preferences based on habitats, and whether this is merely a result of feeding choices occurring among a different set of hosts, or whether it reflects differences in how hosts are distributed across the landscape, the time of day they are present, or other behavioral or physiological changes in hosts or vectors.

Gray catbirds have very low competence for WNV and therefore likely do not contribute to transmission. They comprised a significant proportion of blood meals here but were rarely observed in the avian point counts. Although this could point to a very high level of feeding preference for this species, it may instead reflect the difficulty of detecting them at the time of year or time of day when the point counts were performed. For that reason, we did not calculate forage ratios for this species. Similarly, detection of different mammal species based on camera trap data may differ based on the detectability of different species in different habitats. For instance, in our western sites, there were no camera trap records for eastern cottontails, and although the number of blood meals taken from this species were indeed greater in the eastern sites, there were still a modest number of cottontail blood meals detected in the western sites as well. In other words, there is a level of uncertainty related to detectability in host availability estimates that would propagate to estimates of feeding preferences of mosquitoes.

American goldfinches were in contrast fed on more frequently in the low turf habitats, and it is notable that this species was a significant predictor of WNV infection rates, along with the degree of turf in the environment. A caveat is that the reservoir competence for goldfinches is not known, and the value for house finches was used as a proxy when calculating their competence index. An interpretation is that WNV prevalence is affected strongly by the overall community reservoir competence, which is driven by a number of species, including the American robin, for which the degree of turf grass appears to serve as a proxy, but not for American goldfinches which may explain variation in WNV prevalence in other habitat types. American goldfinches have been shown to have a preference for wet meadows with greater structural diversity, such as open pockets of water and presence of deciduous trees, and more diverse shrub foliage [67].

The abundance of *Cx. pipiens* and *Cx. restuans* was most strongly positively affected by the amount of impervious surface surrounding the collection locations, while the extent of canopy cover showed a negative relationship. An association between developed landcover and *Culex* abundance has previously been noted, although the negative relationship between canopy cover and *Culex* abundance differs from a previous study in central Illinois that found higher rates of oviposition in areas with greater canopy cover [21,68]. The mechanisms involved in the observed pattern could include a correlation between impervious surface and availability of larval development sites such as stormwater catch basins that could promote abundance. While shade and lower temperatures could lower mortality for adult mosquitoes, the increased temperatures associated with increased levels of impervious surface (Fig. S2) may also increase the rate of larval development and lead to an increase in population size. Similarly, increased vapor pressure deficits associated with more urban environments could reduce adult longevity, while promoting blood-feeding activity [69,70]. Further research on how these aspects of the urban environment shape different mosquito traits would be helpful.

Overall, if turf grass can indeed serve as a proxy for the avian community composition that tends to promote WNV infection rates, this offers an opportunity for urban designers, land planners, and residents to consider how different types of urban green space could promote a healthy environment that accounts for exposure to mosquito-borne viruses. There is a need for further research to understand both how generalizable these results are to other urban environments and the potential ramifications of reducing areas with large amounts of turf grass with more diverse habitats, such as urban wetlands or urban forested areas, to help guide urban planning to benefit both urban biodiversity and public health.

## Acknowledgements

We thank all the INHS Medical Entomology Lab field collectors and technicians who provided additional aid with this research project, the field site superintendents who facilitated this work, and Arif Ciloglu for helpful comments. This work was supported by the State of Illinois Used Tire Management and Emergency Public Health funds.

## Notes

### Competing Interest Statement

The authors have declared no competing interest.

