## Supplemental Tables and Figures for "Infection and host-feeding patterns of West Nile virus vectors varies by urban greenspace composition"

**Table S1:** Estimated proportion of total grass/herbaceous ground cover (from US EPA EnviroAtlas Meter-scale Urban Land Cover dataset) that is composed of managed turf grass, based on parcel land use, the percentage of impervious cover (NLCD) within parcel boundary, and parcel area (acres).

<table>
<tr><th rowspan="2">Land Use Classes</th><th rowspan="2">Specific Land Use Types</th><th colspan="3">< 10 % impervious</th><th colspan="2">≥ 10 % impervious</th></tr>
<tr><th>< 10 acres</th><th>10 to 999 acres</th><th>≥ 1000 acres</th><th>< 10 acres</th><th>≥ 10 acres</th></tr>
<tr><td>Residential, Commercial</td><td>all types</td><td rowspan="6">1</td><td rowspan="6">0.7</td><td rowspan="6">0.7</td><td rowspan="6">1</td><td rowspan="6">1</td></tr>
<tr><td>Institutional</td><td>medical, education, religious, cemeteries, other</td></tr>
<tr><td>Industrial</td><td>general, manufacturing / processing, warehousing / distribution, flex</td></tr>
<tr><td>Transportation / Communication / Utility / Waste</td><td>roadway, aircraft, independent auto parking, other</td></tr>
<tr><td>Open Space</td><td>recreation, golf course</td></tr>
<tr><td>Construction</td><td>residential, commercial</td></tr>
<tr><td>Institutional</td><td>gov’t admin & services, prisons, national laboratories</td><td rowspan="7">1</td><td rowspan="7">0.5</td><td rowspan="7">0.3</td><td rowspan="7">1</td><td rowspan="7">0.7</td></tr>
<tr><td>Industrial</td><td>mineral extraction, storage</td></tr>
<tr><td>Transportation / Communication / Utility / Waste</td><td>rail ROW, communication, utility ROW, wastewater treatment, landfill, stormwater mgmt., intermodal, other</td></tr>
<tr><td>Agriculture</td><td>all types</td></tr>
<tr><td>Vacant</td><td>residential, commercial, industrial</td></tr>
<tr><td>Construction</td><td>industrial, other</td></tr>
<tr><td>Water, Non-parcel</td><td>ROW, NEC</td></tr>
<tr><td>Open Space</td><td>conservation, non-public, trail / greenway</td><td rowspan="3">0.7</td><td rowspan="3">0.3</td><td rowspan="3">0.3</td><td rowspan="3">0.7</td><td rowspan="3">0.5</td></tr>
<tr><td>Vacant</td><td>other</td></tr>
<tr><td>Non-parcel</td><td>open space, water</td></tr>
</table>

**Table S2:** Average proportion of female *Culex (Cx.)* spp. specimens collected in CDC light traps and gravid traps lacking sufficient morphologic features to allow identification to species.

| Sample type | Number of samples | Mean proportion of specimens in sample not identified to species ± SE |
|---|---|---|
| Gravid traps | 422 | 0.73 ± 0.05 |
| CDC light traps | 381 | 0.37 ± 0.01 |

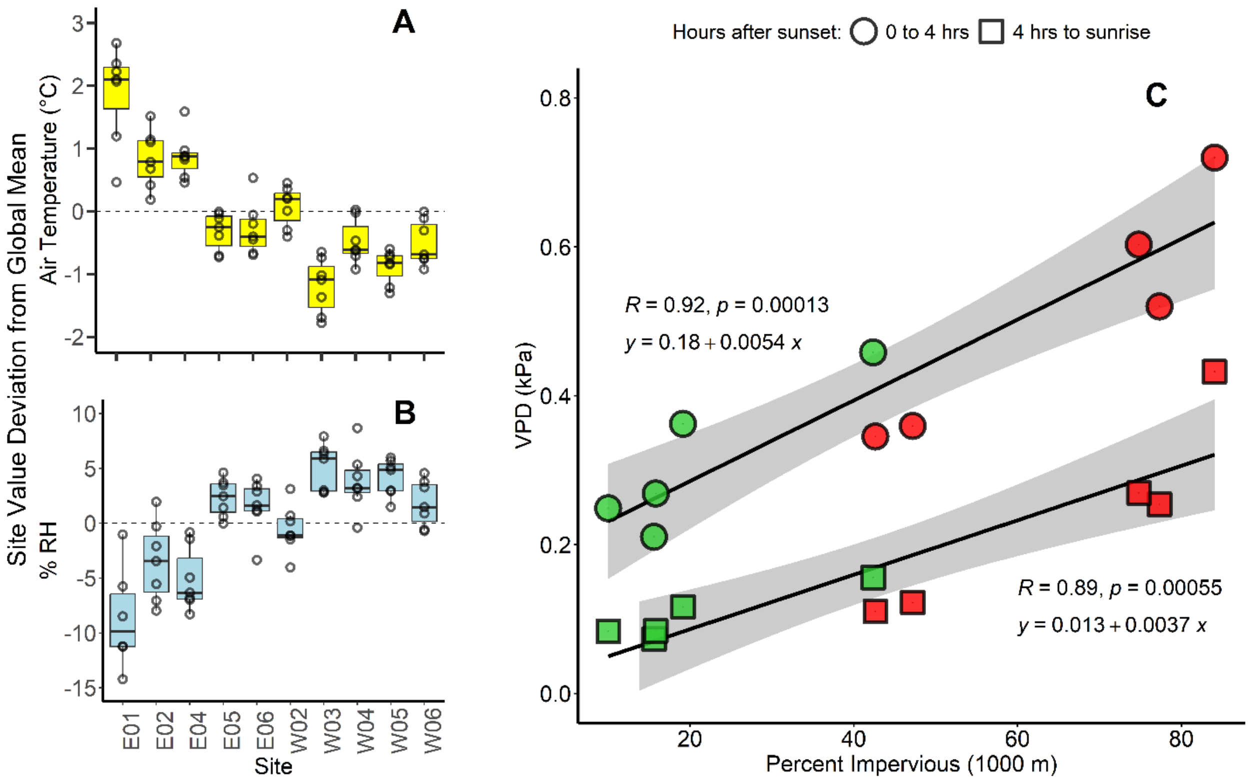


**Fig. S1.** Relative differences in (A) nocturnal air temperature and (B) % relative humidity among sites, and (C) the relationship of percentage impervious surface cover in a 1000 m radius around sites (red = east part of transect, green = west part of transect) to mean vapor pressure deficit during diel periods of expected peak (0 to 4 hours post sunset) and reduced (4 hours post sunset to sunrise) *Culex* spp. flight activity.

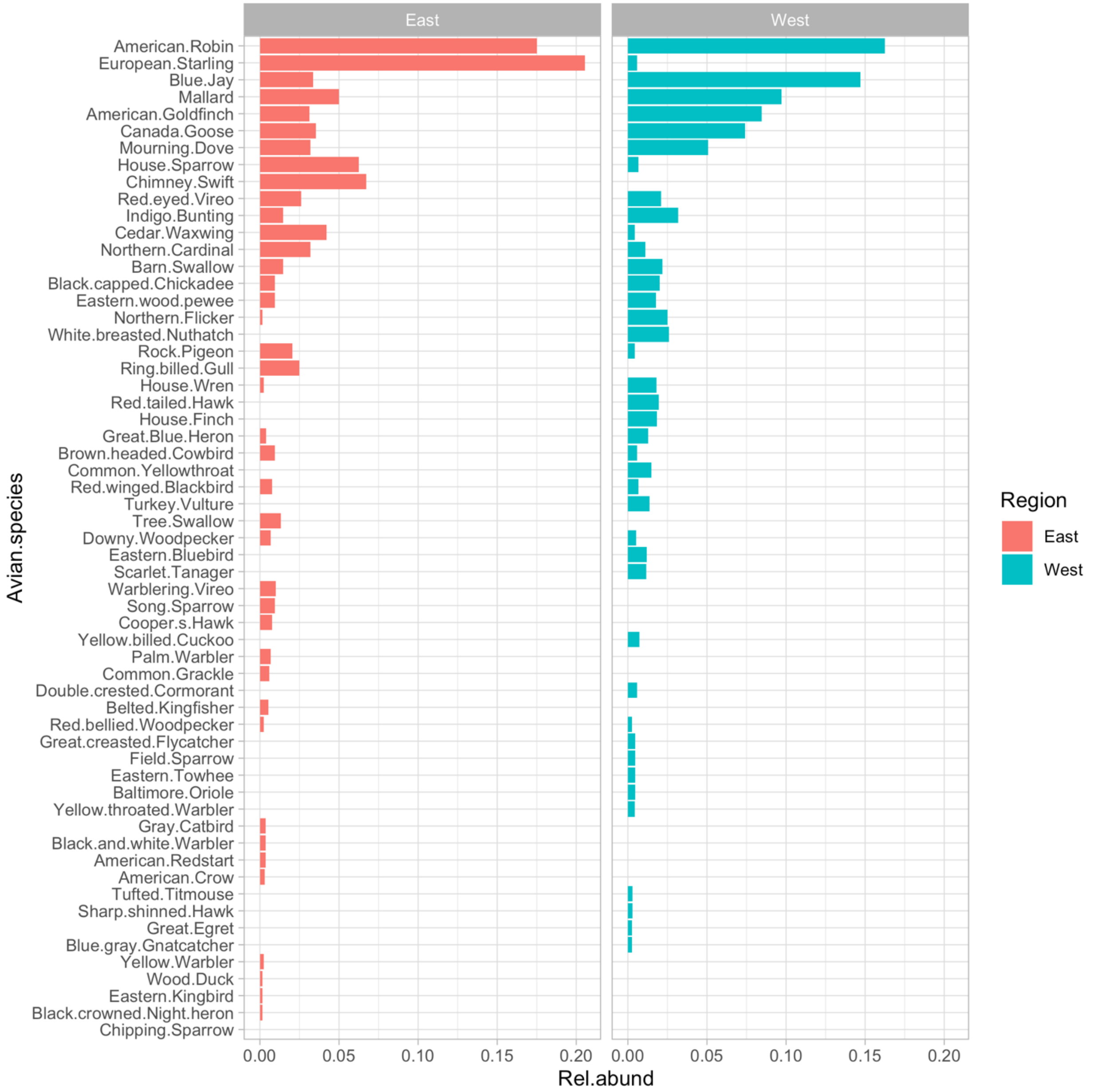


**Fig S2:** Relative abundance of avian species detected over all ten sites grouped by location on the eastern or western side of the transect as determined by point counts.

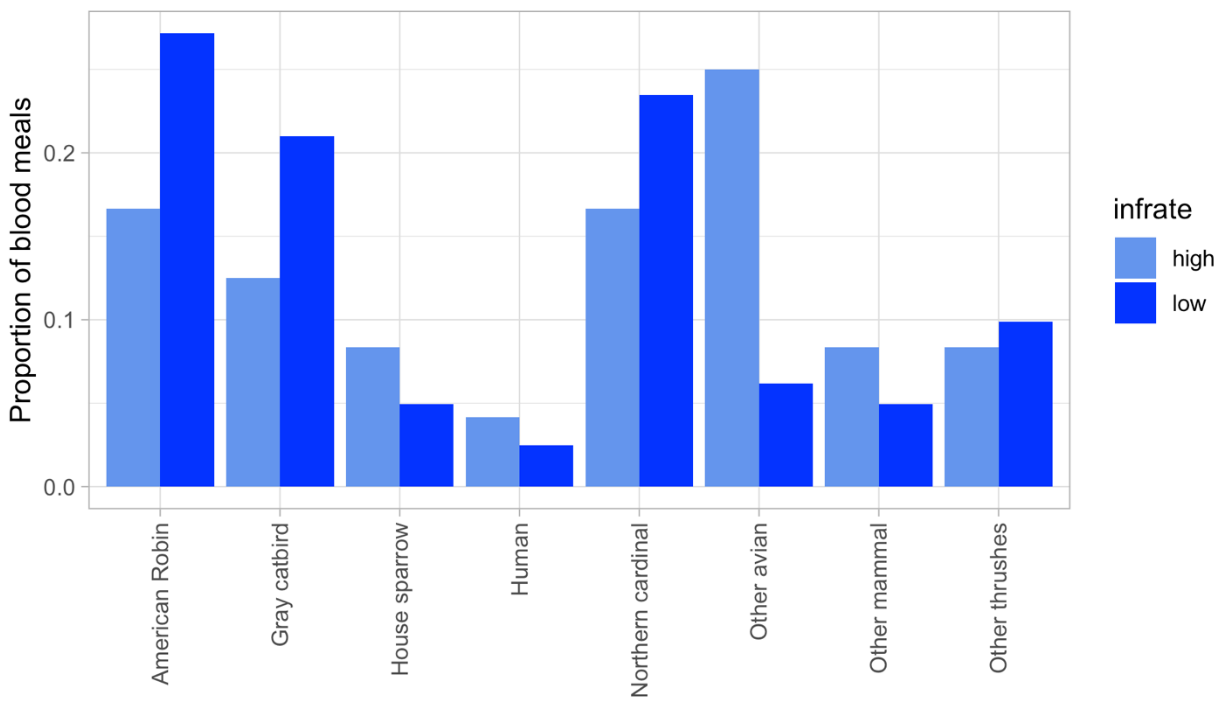


**Fig. S3.** *Cx. pipiens* distribution of blood meals over hosts by sites that had the highest or lowest infection rates.

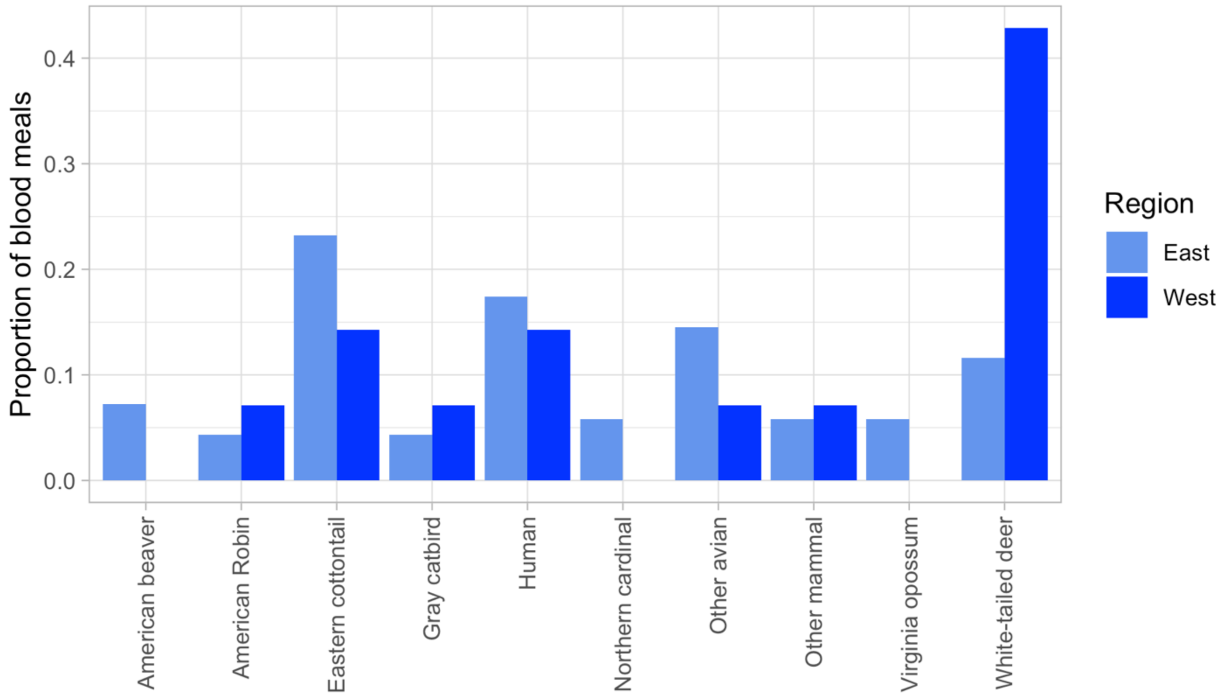


**Fig. S4.** *Cx. salinarius* distribution of blood meals over hosts by sites in the eastern and western regions of the sampling transect.
